# Low cost, high performance processing of single particle cryo-electron microscopy data in the cloud

**DOI:** 10.1101/016451

**Authors:** Michael A. Cianfrocco, Andres E. Leschziner

## Abstract

The advent of a new generation of electron microscopes and direct electron detectors has realized the potential of single particle cryo-electron microscopy (cryo-EM) as a technique to generate high-resolution structures. However, calculating these structures requires high performance computing clusters, a resource that may be limiting to many likely cryo-EM users. To address this limitation and facilitate the spread of cryo-EM, we developed a publicly available ‘off-the-shelf’ computing environment on Amazon’s elastic cloud computing infrastructure. This environment provides users with single particle cryo-EM software packages and the ability to create computing clusters that can range in size from 16 to 480+ CPUs. Importantly, these computing clusters are also cost-effective, as we illustrate here by determining a near-atomic resolution structure of the 80S yeast ribosome for $28.89 USD in ~10 hours.

## Introduction

Cryo-electron microscopy (cryo-EM) has long served as an important tool to provide structural insights into biological samples. Recent advances in cryo-EM data collection and analysis, however, have transformed single particle cryo-EM (Bai et al., 2015; Kuhlbrandt, 2014), allowing it to achieve resolutions better than 5 Å for samples ranging in molecular weight from the 4 MDa eukaryotic ribosome (Bai et al., 2013) to the 170 kDa membrane protein γ-secretase (Lu et al., 2014). These high-resolution structures are the result of a new generation of cameras that detect electrons directly without the need of a scintillator, which results in a dramatic increase in the signal-to-noise ratio relative to CCD cameras, the previous most commonly used device (McMullan et al., 2009). In addition to direct electron detection, the high frame rate of these cameras allows each image to be recorded as a ‘movie’, dividing it into multiple frames. These fractionated images can be used to correct for sample movement during the exposure, further increasing the quality of the cryo-EM images (Campbell et al., 2012; Li et al., 2013; Scheres, 2014).

In addition to these technological developments in the detectors, improvements to computer software packages have played an equally important role in moving cryo-EM into the high-resolution era. Atomic or near-atomic structures have been obtained with software packages such as EMAN2 (Tang et al., 2007), Sparx (Hohn et al., 2007), FREALIGN (Grigorieff, 2007), Spider (Frank et al., 1996), and Relion (Scheres, 2012, 2014). In general, obtaining these structures involved computational approaches that sorted out the data into homogenous classes that could then be refined to high resolution.

While these advances in microscopy and analysis have been essential for the recent breakthroughs in cryo-EM, their implementation is computationally intensive and requires high-performance computing clusters. A recent survey (Scheres, 2014) of high-resolution single particle cryo-EM structures showed that refinement of these structures required processing times > 1,000 CPU-hours (Table 1). Therefore, computational time (i.e. access to high-performance clusters) may represent a significant barrier to determining high-resolution structures by single particle cryo-EM.

**Table 1:** Overview of 3D refinement processing times for recent near-atomic cryo-EM structures.

| **3D refinement processing times<sup>§</sup>** |  |  |  |
| --- | --- | --- | --- |
|  | **Resolution** | **CPU Hours [Months]** | **Amazon EC2 cost** |
| **80S Eukaryotic ribosome<sup>#</sup>** | 4.6 Å | 578 [0.8] | \$12.65<sup>^</sup> |
| **γ-secretase<sup>¶</sup>** | 4.5 Å | 4,660 [6.5] | \$101.94* |
| **β-galactosidase<sup>‡</sup>** | 4.0 Å | 1,160 [1.6] | \$25.38* |
| **Mitochondrial ribosome<sup>†</sup>** | 3.3 Å | 9,330 [13.0] | \$204.09* |

§ Adapted from (Scheres, 2014)

In order to expand the computational resources available to the structural biology community, we investigated the performance of Amazon’s elastic cloud computing (EC2) infrastructure in processing cryo-EM data. We found that the Amazon’s EC2 environment was very amenable to and affordable for the analysis of large cryo-EM datasets; we were able to determine a 4.6 Å structure of the 80S ribosome using a published dataset (Bai et al., 2013) for $28.89 USD in ~10 hours using 128 CPUs. To help others utilize this resource, we compiled an ‘off-the-shelf’ software environment that allows new users to start up a cluster of Amazon CPUs preinstalled with cryo-EM software (Relion, EMAN2, Sparx, FREALIGN, Spider, EMAN, and XMIPP).

## Elastic cloud computing through Amazon Web Services

Amazon Web Services (AWS) is a division of Amazon that offers a variety of cloud-based solutions for website hosting and high-performance computing, amongst other services. Many different types of privately held companies take advantage of Amazon’s computing infrastructure because of its affordability, flexibility, and security. Of note, global biotechnology companies such as Novartis, Bristol-Myers-Squibb, and Pfizer have utilized the computing power of Amazon for scientific data processing. Many academic researchers have also begun to use Amazon’s EC2 resources for analyzing such datasets as super-resolution light microscopy images (Hu et al., 2013), genomics (Krampis et al., 2012; Yazar et al., 2014), and proteomics (Mohammed et al., 2012; Trudgian and Mirzaei, 2012).

The overall workflow starts with users logging into a virtual machine (‘instance’) on AWS (Figure 1). AWS offers a variety of instance types that have been configured for different computing tasks. For example, instances have been optimized for computing performance, GPU-based calculations, or memory-intensive calculations. After logging onto an instance, storage drives are mounted onto the instance, allowing data to be transferred onto the storage drive that can be encrypted for data security (Figure 1).

**Figure 1:**
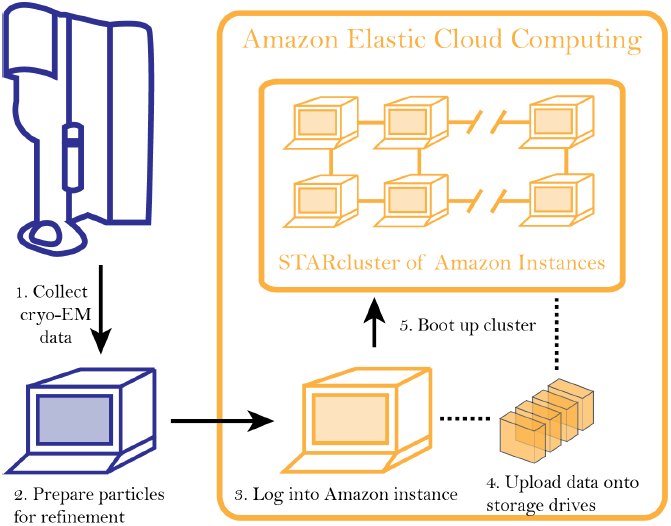
Workflow for analyzing cryo-EM data on Amazon’s cloud computing infrastructure. After collecting cryo-EM data (Step 1), particles are extracted from the micrographs and prepared for further analysis (Step 2). After logging into an ‘instance’ (Step 3), data are uploaded to a storage server (elastic block storage) (Step 4). At this point, a STARcluster can be configured to launch a cluster of 2 – 30 instances that is mounted with the data from the storage volume (Step 5). A detailed protocol can be found at an accompanying Google site: http://goo.gl/AIwZJz.

While users can utilize a single instance for calculations, the maximum number of CPU cores per instance is 18. Therefore, creating a computing cluster on AWS requires additional steps. The Software Tools for Academics and Researchers (STAR) group at Massachusetts Institute of Technology developed a straightforward package that allows users to group individual AWS instances into a cluster. The STARcluster program is a python-based, open source package that automatically creates a cluster preconfigured with the necessary software to manage a computer cluster (Ivica et al., 2009). This package allows users to specify the number of instances to be included in the clusters as well as the instance type. By taking advantage of this tool, private clusters can be built with sizes ranging from 16 to 480 CPUs (Figure 1).

## EM-packages-in-the-Cloud: A pre-configured software environment for single-particle cryo-EM image analysis

We have taken advantage of the ease with which larger numbers of CPUs can be accessed on AWS to compile an off-the-shelf software environment preconfigured with STARcluster and the following cryo-EM software packages: Relion (Scheres, 2012, Scheres 2014), FREALIGN (Grigorieff, 2007), EMAN2 (Tang et al., 2007), Sparx (Hohn et al., 2007), Spider (Frank et al., 1996), EMAN (Ludtke et al., 1999), and XMIPP (Sorzano et al., 2004). We have made this environment publicly available by saving it within Amazon as an ‘Amazon Machine Image’ (AMI), under the name ‘EM-packages-in-the-Cloud.’ In addition to the EM-packages-in-the-Cloud AMI, we also created a second AMI, ‘EM-packages-in-the-Cloud-Node’, which is preconfigured with the necessary STARcluster software required for the cluster nodes. These two publicly available AMIs allows users to boot up a cluster in a few short steps. The protocols describing this can be found on a Google site that is being launched in conjunction with this article: http://goo.gl/AIwZJz. In addition to detailed instructions, the site includes a help forum to facilitate a conversation on cloud computing for single particle cryo-EM.

To test the performance of a cloud-based cluster of Amazon instances, we performed 3D classification and refinement of a previously published 80S *Saccharomyces cerevisiae* ribosome dataset (Bai et al., 2013) on a variety of cluster configurations using Relion. Due to the large memory requirements for Relion’s 3D classification, we ran our performance tests on clusters of high-memory instances — 256 GiB RAM and 16 CPUs per instance, using the ‘r3.8xlarge’ instance-type. Comparison of performance across cluster sizes showed that 256 CPUs had the fastest overall time and the highest speedup relative to a single CPU for both 3D classification and refinement (Figure 2A & 2B). Despite this, cluster sizes of 128 and 64 CPUs were the most cost effective for 3D classification and refinement, respectively, as these were the cluster configurations where the price per speedup increase reached a maximum (Figure 2C). Importantly, the average time required to boot up these STARclusters was ≤ 10 minutes for all cluster sizes (Figure 2D) and, once booted up, the clusters do not have any associated job wait times. Therefore, these tests showed that Amazon’s EC2 infrastructure was amenable to the analysis of single particle cryo-EM data using Relion.

**Figure 2:**
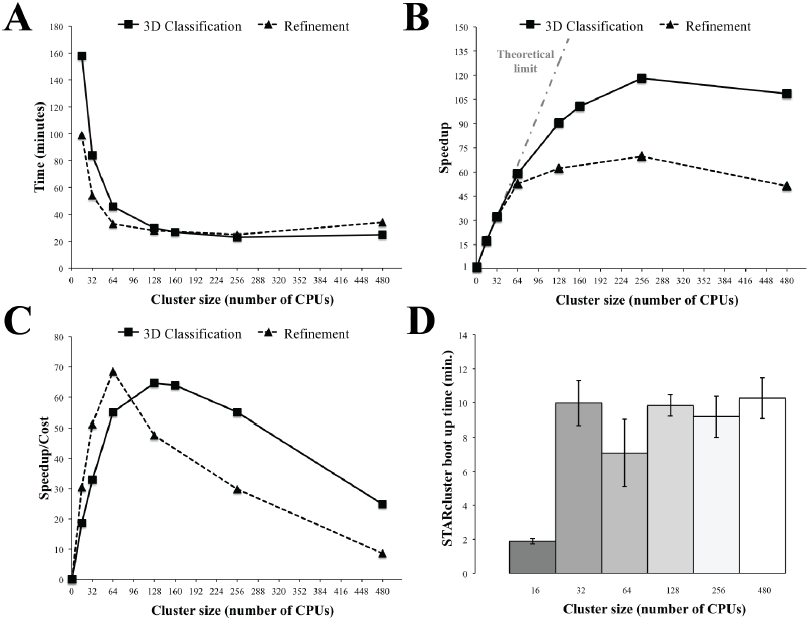
Relion performance on STARcluster configurations of Amazon instances. (A) Processing times (minutes) for Relion to perform 3D Classification or 3D refinement on 80S ribosome dataset. (B) Speedup for each cluster size relative to a single CPU (black line) shown alongside performance estimate for a perfectly parallel cluster using Amdahl’s Law (curve labeled “Theoretical limit”). For cluster sizes ≤ 64 CPUs, the Relion exhibits near-perfect performance on STARcluster configurations, while cluster sizes > 64 show that Relion’s performance reaches a maximum at 256 CPUs for both 3D classification and 3D refinement. (C) Speedup/Cost is plotted against cluster size, where Speedup/Cost is defined as the speedup observed divided by the cost associated with Amazon’s pricing at $0.35/hr/16 CPUs. (D) Average STARcluster boot up time (+/- s.d.) was measured for clusters of increasing size (n = 5).

Further analysis of the 80S ribosome dataset on a 128 CPU cluster (8 x 16 CPUs; using the r3.8xlarge instance) highlighted the cost-effectiveness of Amazon’s EC2 infrastructure. After 3D classification, a homogenous group of particles was refined to an overall resolution of 4.6 Å (Figure 3A - C). This structure, which involved 3D classification and refinement, required a total of ~10 hours on a 128 CPU cluster. By bidding on unused instances (‘spot instances’), each 16 CPU instance cost $0.35/hr instead of $2.80/hr, saving 87.5% per instance. At this price, the 4.6 Å ribosome structure had an overall cost of $28.89 (8 instances x $0.35/hr x 10.3 hrs). Thus, even though obtaining this structure required 1,321 total CPU-hours, Amazon’s EC2 computing infrastructure provided the resources to calculate it to near-atomic resolution within ~10 hours, with a total cost of $28.89.

**Figure 3:**
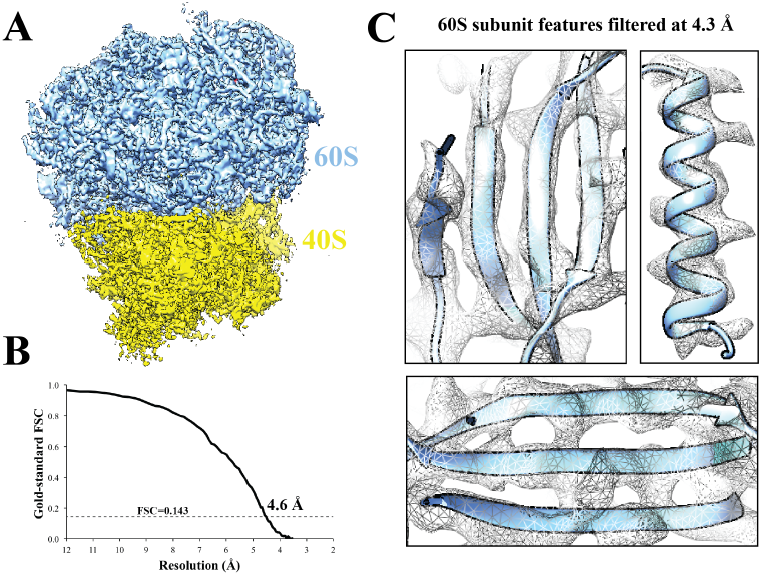
Cryo-EM structure of 80S ribosome at an overall resolution of 4.6 Å. (A) Overall view of 80S reconstruction filtered to 4.6 Å while applying a negative B-factor of -116 Å^2^. (B) Gold standard FSC curve. (C) Selected regions from the 60S subunit filtered at 4.3 Å. Cryo-EM maps were visualized with UCSF Chimera (Pettersen et al., 2004). **Source data:** A detailed description of each step of the image processing is available at https://github.com/mcianfrocco/Cianfrocco-and-Leschziner-2014-EMCloudProcessing/wiki. Associated computing scripts and data files have been uploaded to Github (https://github.com/mcianfrocco/Cianfrocco-and-Leschziner-2014-EMCloudProcessing) and Dryad Digital Repository (http://datadryad.org/review?doi=doi:10.5061/dryad.9mb54) (Cianfrocco and Leschziner), respectively.

## Cloud computing as a tool to facilitate high-resolution cryo-EM

Recent advances in single particle cryo-EM have drawn the interest of the broader scientific community. In addition to technical advances in electron optics, the new direct electron detectors and data analysis software have dramatically improved the resolutions that can be achieved for a variety of structural targets. In contrast to the other high-resolution techniques (X-ray crystallography, NMR), structure determination by cryo-EM is extremely computationally intensive. The publicly available ‘EM-packages-in-the-Cloud’ environment we have presented and characterized here will help remove some of the limitations imposed by these computational requirements. Comparisons of predicted refinement costs for published structures showed that Amazon’s cloud computing network is a powerful, cost-effective tool for analyzing cryo-EM data (Table 1).

We believe that cloud-based approaches will impact the future of cryo-EM image processing in two fronts: 1) new cryo-EM users / laboratories across the globe will have immediate access to a high performance cluster, and 2) existing labs will have a dramatic increase in productivity. As the number of laboratories using cryo-EM increases, and as existing laboratories begin to pursue high-resolution cryo-EM, gaining immediate access to a high performance cluster may become difficult. For instance, while there are government-funded high performance clusters in the United States (e.g. XSEDE STAMPEDE), it may take up to a month for a user application to be reviewed (Rogelio Hernandez-Lopez, personal communication). Assuming that the application is approved, these clusters may not have appropriate software installed, which further delays cluster usage. Finally, the user will have a set limit for the number of CPU hours available per project, requiring a new application to be submitted to access the cluster again. All of these problems are circumvented by using Amazon’s EC2 infrastructure, which provides immediate, cost-effective access to hundreds of CPUs for users around the globe.

While many research laboratories have previously bought and maintained private clusters (200 – 500 CPUs), we believe that Amazon’s EC2 infrastructure will provide more processing per lab member at reduced costs. The price of a high performance private cluster with 320 CPUs costs ~$50,000, which does not include costs associated with housing and maintaining the cluster. Even after setting up the private cluster, it will likely be shared across a laboratory of 5 – 10 people, reducing the effective number of CPUs available per lab member. Therefore, considering the price of using Amazon’s EC2 infrastructure, laboratories will be able to have 300+ CPUs per lab member for much less than it would cost to buy and maintain a private cluster.

By minimizing computational time and increasing global accessibility, high-performance cloud computing can help usher in the era when high-resolution cryo-EM becomes a routine structural biology tool.

## Materials and Methods

### Setting up a cluster on Amazon EC2 with spot instances

In order to minimize costs, STARclusters were assembled from ‘spot instances,’ which are unused instances that are open to a bidding process in order to be reserved. The spot instances are different from ‘on-demand’ instances: on-demand instances provide users with guaranteed access while spot instances are reserved until there is a higher bid, at which point the user is logged out of the spot instance. For example, while the on-demand rate for r3.8xlarge instances is $2.80/hr, the minimum bidding price for a spot instance of r3.8xlarge is $0.25/hr. This means that a user could reserve a spot instance for as little $0.25/hr, as long as no other users bid >$0.25/hr.

Throughout our experience on Amazon’s EC2 (2 months), we found that the average spot instance price for r3.8xlarge was $0.25 - $0.28/hr. This meant that bidding $0.28 - $0.30/hr provided near-guaranteed access to the spot instances in the US East Region (N. Virginia), where we have only been kicked off the instances twice. A protocol describing the necessary steps to set up a cluster has been detailed on an accompanying website: http://goo.gl/AIwZJz.

### CPUs vs. vCPUs

In selecting an instance type, new users should be aware of the differences between CPUs and vCPUs on Amazon’s EC2 network. Namely, that there are two vCPUs per physical CPU on Amazon. This means that while r3.8xlarge instances have 32 vCPUs, there are actually only 16 physical CPU cores in each instance, with each CPU having two hyperthreads. Practically, this means that Amazon’s instances have higher performance than a 16 CPU machine and less performance than a 32 CPU machine. To account for this difference, all numbers reported here were CPU numbers that were converted from vCPUs.

### Image processing

Micrographs from the 80S *Saccharomyces cerevisiae* ribosome dataset (Bai et al., 2013) were downloaded from the EMPIAR database for electron microscopy data (EMPIAR 10002). The SWARM feature of EMAN2 (Tang et al., 2007) was used to pick particles semi-automatically. Micrograph defocus was estimated using CTFFIND3 (Mindell and Grigorieff, 2003). The resulting particle coordinates and defocus information were used for particle extraction by Relion-v1.3 (Scheres, 2012, 2014). The particle stacks and associated data files were then uploaded to an elastic block storage volume on Amazon’s EC2 processing environment at a speed of 10 MB/sec (24 minutes total upload time).

Initial 3D classification was performed on 62,022 80S Ribosome particles (1.77 Å/pixel). These were classified into 4 groups (T=4) for 13 iterations using a ribosome map downloaded from the Electron Microscopy Data Bank (EMDB-1780) that was low pass filtered to 60 Å. Further 3D classification using a local search of 10° and an angular sampling of 1.8° continued for 13 iterations. At this point, two classes were identified as belonging to the same structural state and were selected for high-resolution refinement (32,533 particles). Refinement of these selected particles continued for 31 iterations using *3D auto-refine* in Relion. The final resolution was determined to be 4.6 Å using *Post process* in Relion, applying a mask to the merged half volumes and a negative B-factor of -116 Å^2^.

### Performance Analysis

80S ribosome data were reanalyzed on clusters of increasing size using both 3D classification and 3D refinement. The time points collected involved running 3D classification for 2 rounds and 3D refinement for 6 rounds, using the same number of particles and box sizes listed above: 62,022 particles for classification and 32,533 particles for refinement with box sizes of 240 x 240 pixels. The Relion commands were identical to the commands used above and the calculations were terminated after the specified iteration.

From these time points, the speedup of each cluster size was calculated relative to a single CPU. Speedup (*S*) was calculated as

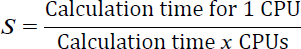

The measured speedup values were then compared to the speedup expected for a perfectly parallel algorithm (***P*** = 1) using Amdahl’s law (Amdahl, 1967):

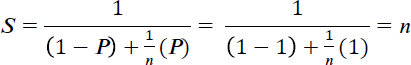

Where ***P*** is the fraction of an algorithm that is parallel and *n* is the number of processors. The calculation time for 3D classification on a single CPU were calculated by using 1 CPU on a 16 CPU r3.8xlarge instance. For calculating a 3D refinement on a single CPU, (or two vCPUs), the refinement was run on 4 vCPUs and then converted to a single CPU (or two vCPUs) by multiplying the calculation time by 2. For cost analysis, the measured speedup was divided by the cost to run the job on spot instances of r3.8xlarge at a price of $0.35/hr. Cluster boot up times were calculated from the elapsed time between submitting the STARcluster command and the STARcluster fully booting up.

## Data Accession Information

Further information regarding ‘EM-Packages-in-the-Cloud’ can be found at an associated Google Site: http://goo.gl/AIwZJz. The final 80S yeast ribosome structure at 4.6 Å has been submitted to the EM Databank as EMDB 2858. A detailed description of each step of the image processing is available at https://github.com/mcianfrocco/Cianfrocco-and-Leschziner-2014-EMCloudProcessing/wiki. Associated computing scripts and data files have been uploaded to Github (https://github.com/mcianfrocco/Cianfrocco-and-Leschziner-2014-EMCloudProcessing) and Dryad Digital Repository (http://datadryad.org/review?doi=doi:10.5061/dryad.9mb54) (Cianfrocco and Leschziner), respectively.

## Acknowledgements

We would like to thank all the members of the Leschziner and Reck-Peterson labs for critical discussions. We would like to especially thank Rogelio Hernandez-Lopez, Anthony Roberts, and Daniel Cianfrocco for critical feedback on the development of this Amazon computing environment. We also would like to thank the Structural Biology Consortium (SBGrid) for pricing information on cluster and file server sizes. M.A.C. is an HHMI fellow of the Damon Runyon Cancer Research Foundation and A.E.L is supported by NIH/NIGMS (R01 GM107214 and R01 GM092895A).

### Competing financial interests statement

The authors do not have any competing financial interests.

